# Selective breeding and production strategies to support snapper farming in the warming waters of New Zealand’s South Island

**DOI:** 10.1101/2025.06.13.659617

**Authors:** Georgia Samuels, Flavio Ribeiro, David Ashton, Sharon Ford, Joshua Fantham, Julie Blommaert, Damian Moran, Maren Wellenreuther

## Abstract

Diversifying aquaculture species is essential for building resilience in the face of climate change, particularly as warming oceans challenge existing production systems. In New Zealand’s Marlborough Sounds, rising sea temperatures are making salmon farming increasingly difficult, highlighting the need for climate-adapted alternatives. This study evaluated the aquaculture performance of selectively bred F_4_ versus unselected F_1_ Australasian snapper (*Chrysophrys auratus*) across two rearing environments: sea pens in the Marlborough Sounds and a land-based system in Nelson. Approximately 1,000 F_4_ and 1,000 F_1_ snapper were reared from 4 to 30 months of age in each system. At 30 months, selectively bred snapper showed improved growth-body length increased by 1.7% (land-based) and 4.8% (ocean-based), and body weight by 9.8% and 14.2%, respectively compared with F_1_ snapper. Survival was also significantly higher, with selected snapper outperforming controls by 84.2% in the land-based and 60.8% in the ocean-based system. Mortality peaked in the first winter across both systems, with size-selective patterns in sea pens informing minimum stocking sizes. These findings offer important insights for refining husbandry and selective breeding practices. They are not only relevant for New Zealand but also for global aquaculture sectors seeking robust species suited to changing marine environments.

## Introduction

Aquaculture, the cultivation of aquatic organisms such as finfish, shellfish, and seaweed, is the fastest-growing food production sector (FAO 2022), currently supplying half the world’s seafood for human consumption (Boyd et al. 2022). As the global population continues to grow, the demand for seafood is projected to rise significantly (Smith et al. 2010; United Nations 2015; De Meester et al. 2024). To meet this growing demand, species diversification in aquaculture has become increasingly crucial. The aquaculture diversification towards native species not only reduces the reliance on wild fisheries but also provides market resilience by means of a constant supply of seafood product to a market that is otherwise seasonal.

Species diversification also expands the range of farmed species with varying environmental and biological requirements, enabling more efficient use of available marine farming areas. Furthermore, species with a wider range of abiotic tolerance, such as variations in temperature tolerance, are ideal candidates for diversification as they can help to optimize the geographical use of grow-out space (Teletchea 2021; Cai et al. 2023).

Climate change is putting an urgent pressure on the aquaculture industry to transit to species that are more tolerant to environment changes. The success and productivity of ocean-based aquaculture depends closely on water temperatures. Rising temperatures and environmental shifts could therefore threaten some of the industry’s ability to sustain growth and maintain fish production in the future (Yadav et al. 2024). This is particularly true for the aquaculture of temperate species, which depends on cold water. Climate change impacts are already disrupting established farming systems around the globe, as seen in Tasmania (Meng et al. 2022) and Norway (Ytteborg et al. 2023; Falconer et al. 2024), and again, this disruption is predicted to increase with ongoing climate warming. The resulting rising ocean temperatures are forecasted to trigger increasing and extended marine heatwaves and associated fluctuating oxygen levels (Law et al. 2018). These events are placing significant stress on many farmed species, leading to higher mortality rates and reduced productivity. Action is needed to prepare the industry for the coming challenges and consequently, it is crucial to introduce new species that can better withstand changing environmental conditions in areas where production of contemporary species is no longer viable.

New Zealand’s ocean-based finfish aquaculture industry is currently dependent on a single species—Chinook salmon (*Oncorhynchus tshawytscha*)— which is farmed exclusively in the South Island. However, the viability of salmon farming is increasingly threatened by changing climatic conditions, particularly in the Top of the South Island, where salmon farms in the Marlborough Sounds have experienced reduced production because of a general warming of summer water temperatures, as well as increasingly frequent marine heatwaves. Notable mortality events associated with marine heatwaves have occurred in 2021 and 2022 (Cook et al. 2024), and high losses were also reported in 2025 (De Boni 2025), highlighting the vulnerability of salmon to climate change. As salmon farming looks to move to cooler waters in the open ocean or further south to adapt to these changing conditions, there are opportunities to develop other species more suited to the future marine climate of the Marlborough Region. This study presents the first evaluation of hatchery-reared Australasian snapper (*Chrysophrys auratus*) on-grown in a net pen in the Marlborough Sounds, with the goal of identifying the key biological challenges that would need to be addressed before proceeding further towards commercial-scale trials.

The Australasian snapper (Sparidae: *Chrysophrys auratus*, also known as tāmure by the indigenous Māori people of New Zealand, hereafter referred to as snapper) is a promising species for aquaculture diversification in New Zealand for several reasons: it has a wide latitudinal distribution (Parsons et al. 2014; Papa et al. 2020); an existing market profile with reasonably good prices ($NZ11 per kg whole fish) from a modest and quota-limited wild-capture sector (∼7000 tonnes) (The economic contribution of commercial fishing 2022); and it is a member of the Sparidae family for which there are well-developed and transferrable aquaculture practices for related species, such as the Mediterranean seabream (*Sparus aurata*) and Japanese red seabream (*Pagrus major*). An understanding of Australasian snapper for aquaculture has been progressively developed over the past 20 or so years. In New Zealand, snapper is predominantly found in the North Island and the northern part of the South Island (Crossland 1981a, 1981b), with the southern end of its distribution range being the southern South Island (Graham 1953), where temperatures can go below 10°C. Like many other Sparidae species, they experience a broad range of natural temperatures (Francis 1996, 2001; Wellenreuther et al. 2019; Moran et al. 2023). For instance, their range in Australia extends as far north as Mackay, where sea surface temperatures (SSTs) can reach up to 30°C (Ferrell 1993). Given snapper has a wide range of thermotolerance, the species could facilitate a climate-resilient aquaculture transition in New Zealand and be farmed in both South and North Islands, under a range of temperature profiles. In Australia, studies have addressed topics such as nutrition (Booth MA et al. 2004; Booth MA et al. 2007; Booth M, Tucker, et al. 2008), harvested fish colour (Booth MA et al. 2004; Doolan et al. 2007) and production in saline inland waters (Fielder et al. 2001). In New Zealand, the focus has been improving the production cycle and on selective breeding to generate high-performing fish and genomic tools for aquaculture, resulting in an elite snapper line with superior growth, survival and feed conversion ratio (FCR) (Moran et al. 2023; Samuels et al. 2024). Snapper has a wide distribution across New Zealand and Australia, inhabiting nearly all inshore environments down to depths of 200 m (Leach 2007; Parsons et al. 2014). To date all published information in New Zealand has relied on tank-based populations; however, this type of data cannot directly address questions about the farming potential of Australasian snapper aquaculture in the Marlborough region, as there is limited transferability of fish performance data between tank-based studies and sea pens.

We sought to benchmark the growth and survival of a selectively bred F_4_ cohort of snapper against a non-selected F_1_ cohort in a remotely operated sea pen in the Marlborough Sounds, while also keeping populations of each cohort in tanks at a land-based facility in Nelson to enable the collection of higher-resolution weight gain and mortality data. A novel aspect of this study was the use of image-based re-identification of individual fish over time, to disentangle the correlated traits of growth and survival in selectively bred and wild-type fish. Size-selective mortality has a significant effect on structuring populations of many fish species, particularly during early life stages when small fish with limited energetic stores encounter adverse conditions (Rosenberg and Haugen 1982; Sogard 1997; Folkvord et al. 2009; Garrido et al. 2015; Le Pape and Bonhommeau 2015; Peres et al. 2022).

Understanding the relationship between fish body weight at the time they are transferred into a sea pen and their subsequent growth and survival performance across seasons is critical to optimising aquaculture production strategies, and if this information can be refined to individual-level data, it can help inform breeding programmes and size-grading thresholds for juvenile fish destined for sea pen on-growing.

## Materials and Methods

### Ethics Statement

This study was approved under animal ethics application AEC-2021-PFR-05 by the Animal Ethics Committee at the Nelson Marlborough Institute of Technology (NMIT), and operations approved under fish-farm license FW208.03.

### Land-based and ocean-based production settings

Land-based on-growing of Australasian snapper (*Chrysophrys auratus*) was conducted at Plant & Food Research’s (PFR) Nelson Research Facility, located in Nelson, New Zealand (41.2985° S, 173.2441° E) (Figure 1A). This facility is equipped with a flow-through system, with water drawn from the Nelson Haven from an engineered aquifer in the intertidal zone filled with various hard substrates that provide solids filtration. Abstracted filtered water is then pumped through the facility in a reticulation system that delivers the water into the fish tanks. Fish tanks are fitted with air diffusers and ceramic oxygen stones to further assist with maintaining water quality and allow exchange of gases. Environmental parameters of the tanks, such as water temperature, were recorded daily and Dissolved Oxygen (DO) was monitored daily around feeding events.

**Figure 1.**
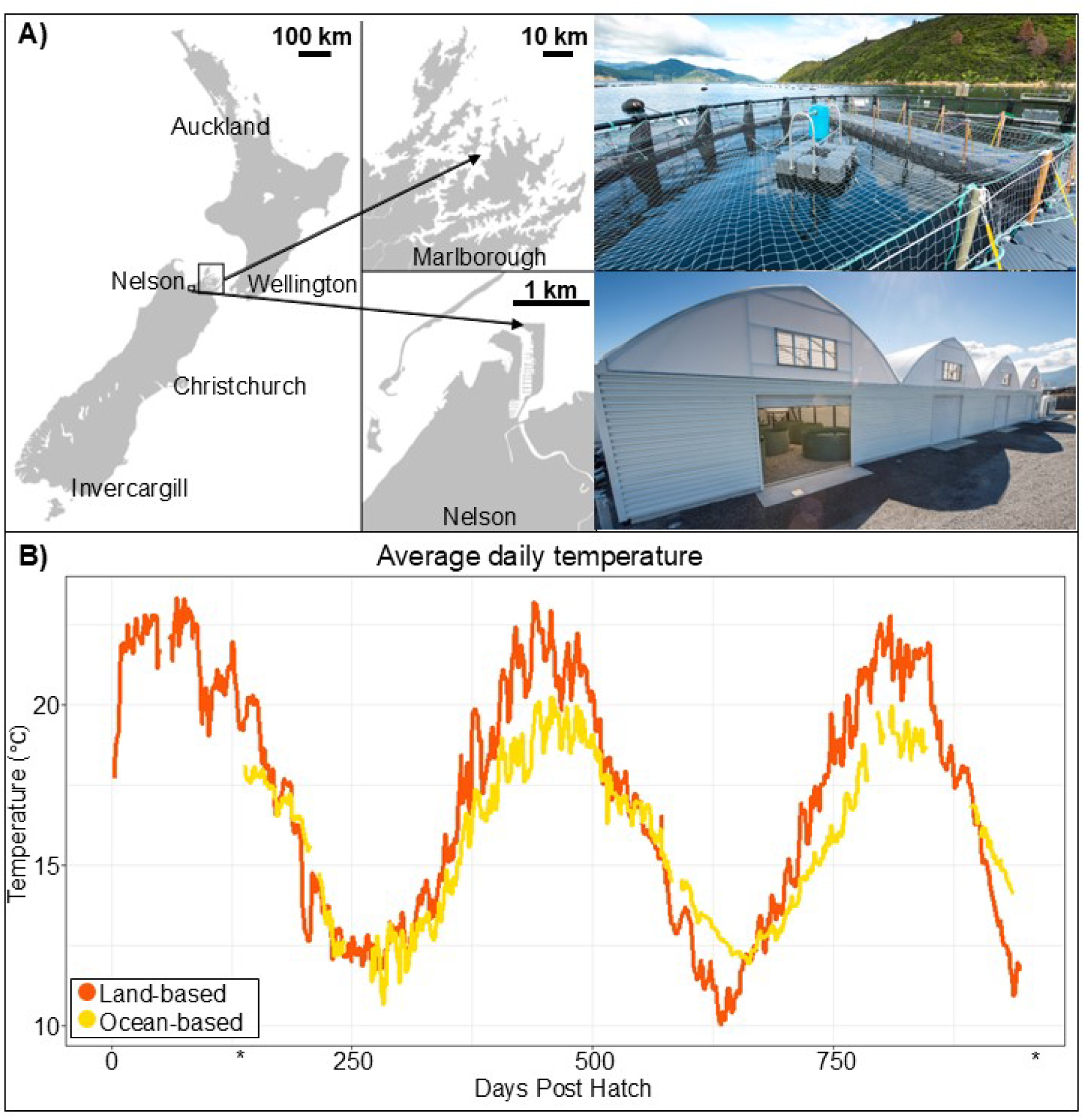
Panel A depicts a map of New Zealand highlighting the production locations, Nelson (land-based) and Marlborough (ocean-based), where Australasian snapper (*Chrysophrys auratus*) were grown in this study. Additionally, this shows the land-based and ocean-based systems next to each location. Panel B displays the temperature profile coloured by production setting, with land-based system shown in orange and ocean-based in yellow. The temperature is shown from incubation of eggs until completion of the study. An asterisk marks 4 and 30 months of age, the start and end points for this comparison.

Ocean-based on-growing of snapper was conducted at a sea pen located in Beatrix Bay, Marlborough Sounds, New Zealand (41.1173° S, 174.0706° E) (Figure 1A). The site, leased from Ngāi Tahu Seafoods, consisted of a 19-meter diameter floating ring pen with a 15-meter deep KikkoNet (Maccaferri Australasia Ltd, Melbourne Australia) mesh enclosure. The main pen was subdivided into four equal sub-pens, each constructed from knotless nylon mesh with a 25 mm bar length, and measuring 6 × 6 × 6 meters (∼216 m³ water volume per sub-pen). Pontoon walkways provided access between sub-pens. The pen system was powered by solar panels and 12 V batteries, which supported an Arvo-Tec (Arvo-Tec Oy, Huutokoski, Finland) automatic feeding system, Adroit (Adroit Ltd, Auckland, New Zealand) environmental monitoring sensors (temperature, DO, pH, turbidity, salinity, redox potential, and conductivity) and SnapIT underwater monitoring and transmitting cameras (SnapCore Ltd, Nelson, New Zealand). The snapper cohort was stocked into one of the four sub-pens for the duration of the trial.

### Production of F_1_ and F_4_ cohorts

F_4_ and F_1_ snapper cohorts were produced in November 2021 using third generation selectively bred broodstock (n = 201) and wild-caught broodstock (n = 61) sourced from the Marlborough Sounds and Tasman Bay, respectively (Samuels et al. 2024). Fertilized eggs from each group were incubated for five days, yielding approximately 17,482 and 22,919 weaned post-larvae at 40 days post hatch (DPH) in the F_4_ and F_1_ cohort, respectively.

### On-growing conditions

On 1 March 2022, 4-month-old snapper (Samuels et al. 2024) were phenotyped (described below) and size graded to remove individuals less than 70 mm fork length (a requirement based on the sea pen mesh size). From the size-graded populations, snapper juveniles were randomly split into four cohorts as follows: Cohort 1: 1204 F_4_ snapper; cohort 2: 1159 F_1_ snapper; cohort 3: 1056 F_4_ snapper; cohort 4: 1056 F_1_ snapper. While cohorts 1 and 2 were retained in the land-based system in two 5,000-L tanks (one for each cohort), cohorts 3 and 4 were transported to the ocean-based system.

For the land-based cohorts, a maximum stocking density of 15 kg m^-3^ was maintained for the duration of the study. Density was assessed at each sampling event and a number of snapper were removed from the tank either to maintain the stocking density at similar rates in between the F_4_ and F_1_ cohorts, or to keep the stocking density within acceptable parameters for water quality and preserve fish welfare. Cohorts 3 and 4 were transported to the sea pen by road using a portable live-fish transport system placed on a truck, which was then transferred to a vessel for the final part of the trip, with sea water exchange while on water. Dissolved oxygen and temperature were monitored throughout the four hours of transportation. On arrival, F_4_ and F_1_ snapper cohorts were transferred into the same sub-pen (initial stocking density of 0.3 kg m^-3^) and remained there for 26 months (final stocking density of 1.9 kg m^-3^).

All four cohorts were fed the same commercial diets. Skretting Nutra RC (2.3-mm to 3.0-mm) was provided from 4 to 12 months of age, while Otohime (4-mm) was offered afterwards until the end of the comparison. For the land-based cohorts, feeding was done by hand, three times a day, to apparent satiation, while for the ocean-based cohorts, food was stored in a 100-L hopper connected to a rotating drum feeder system (Figure 1A), which automatically delivered food at regular intervals during daylight hours at a rate of approximately 1.5% body weight per day.

Land-based cohorts were monitored daily, and ocean-based cohorts every two to four days for mortalities. At the conclusion of the comparison, all remaining ocean-based stock were live-transported back to the land-based system in Nelson using the system previously described. All individuals from both the land-based and ocean-based cohorts were then counted and assessed simultaneously to finalize the study.

### Trait measurements

Initial phenotyping of all individuals was conducted at 4 months of age, where fish in both the land-based and ocean-based cohorts were manually measured for fork length (*L_F_*, mm) and body weight (BW, g) for growth monitoring. Furthermore, associated high-quality images of the left side of the snapper were captured for image analysis (described later). Growth performance data were periodically monitored at 11, 14, 18, and 24 months through a subsample of a minimum 10% of the population. At 30 months of age, all individuals were measured to conclude the study (Figure 2 A–B and Table 1). To generate detailed growth curves, the land-based cohort was sub-sampled at an additional eight time points (5, 7, 12, 15, 16, 23, 26, and 27 months of age). Prior to measurement, snapper were anaesthetised using AQUI-S® (Aqui-S New Zealand Ltd, Lower Hutt: 15–20 ppm), a dose which resulted in a loss of equilibrium and no reaction to net capture. In the land-based cohorts, sub-samples of anesthetised snapper were captured from tanks using dip nets and transferred to a measurement station until the target sample number was reached. For the ocean-based cohorts, the stock was crowded by raising the base of the net and then lining with a purpose-designed tarpaulin bath dosed with AQUI-S® (8–10 ppm) to sedate the snapper. From here, snapper were dip-netted and counted into a 680-L bin until the target sample number was reached. Individuals were then further anaesthetised using AQUI-S® (15–20 ppm) and processed through a measurement station for size and image recording. On 8 May 2024, all remaining ocean-based stock were retrieved from the sea pen (850 individuals at 30 months old) and transferred back into the land-based system for final phenotyping.

**Figure 2.**
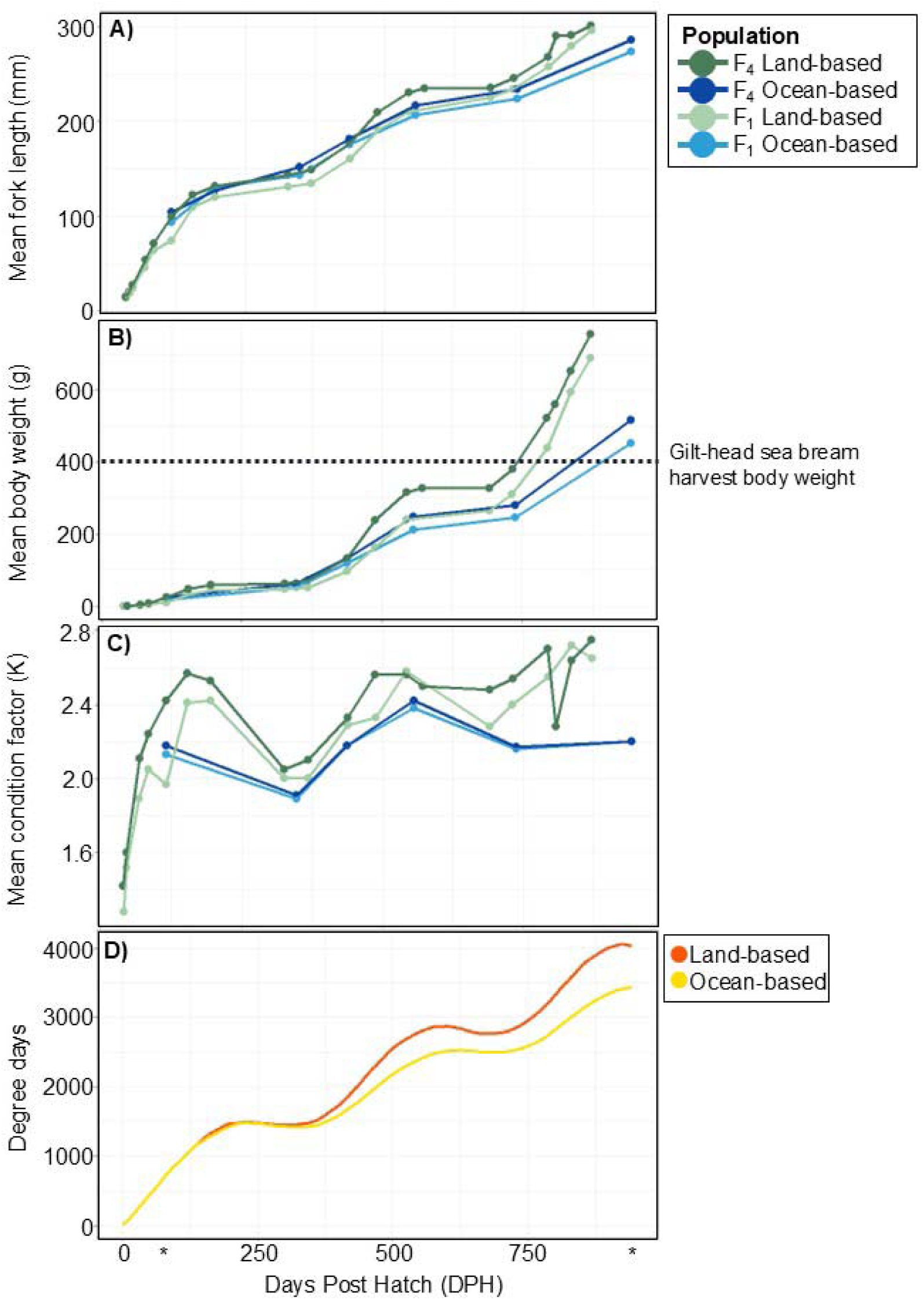
Mean fork length (panel A), mean body weight (panel B), mean condition factor (K) (panel C) of F_4_ and F_1_ Australasian snapper (*Chrysophrys auratus*), and degree days (panel D) for cohorts in land-based and ocean-based systems. Graphs are from egg incubation to end of the study. Asterisks mark 4 and 30 months of age, the start and end points for the direct comparison. Additionally, panel B highlights the harvest body weight (g) for the gilt-head sea bream (*Sparus aurata*) (Mhalhel et al. 2023), the sister species to the Australasian snapper. Graphs have each been coloured by cohort.

**Table 1.**
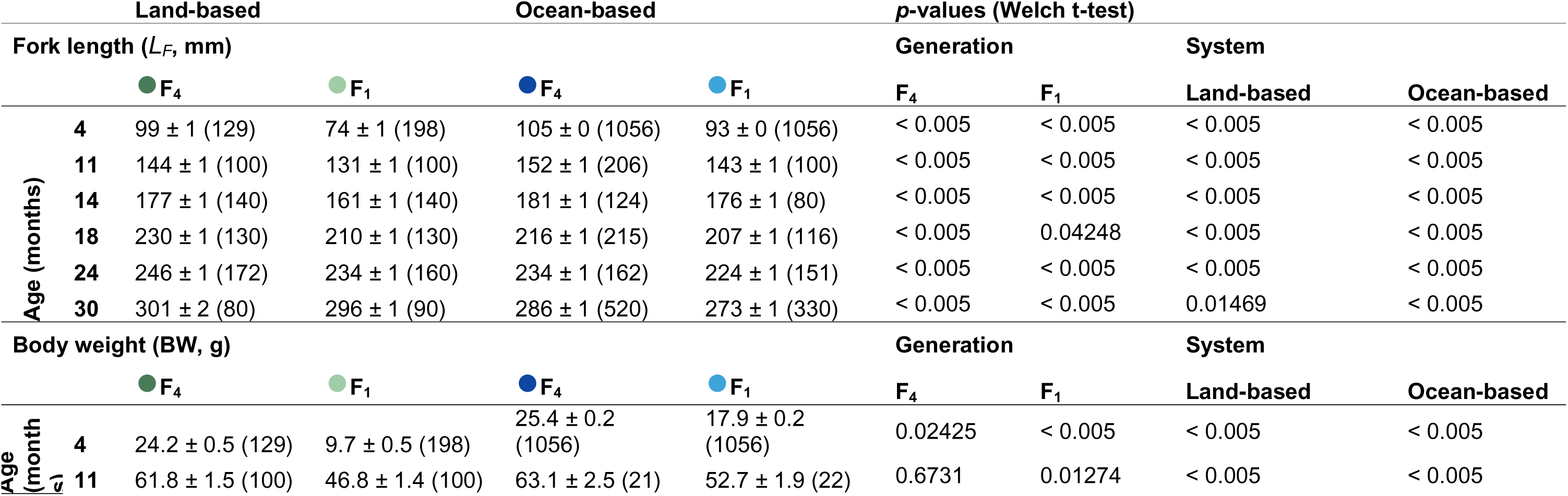
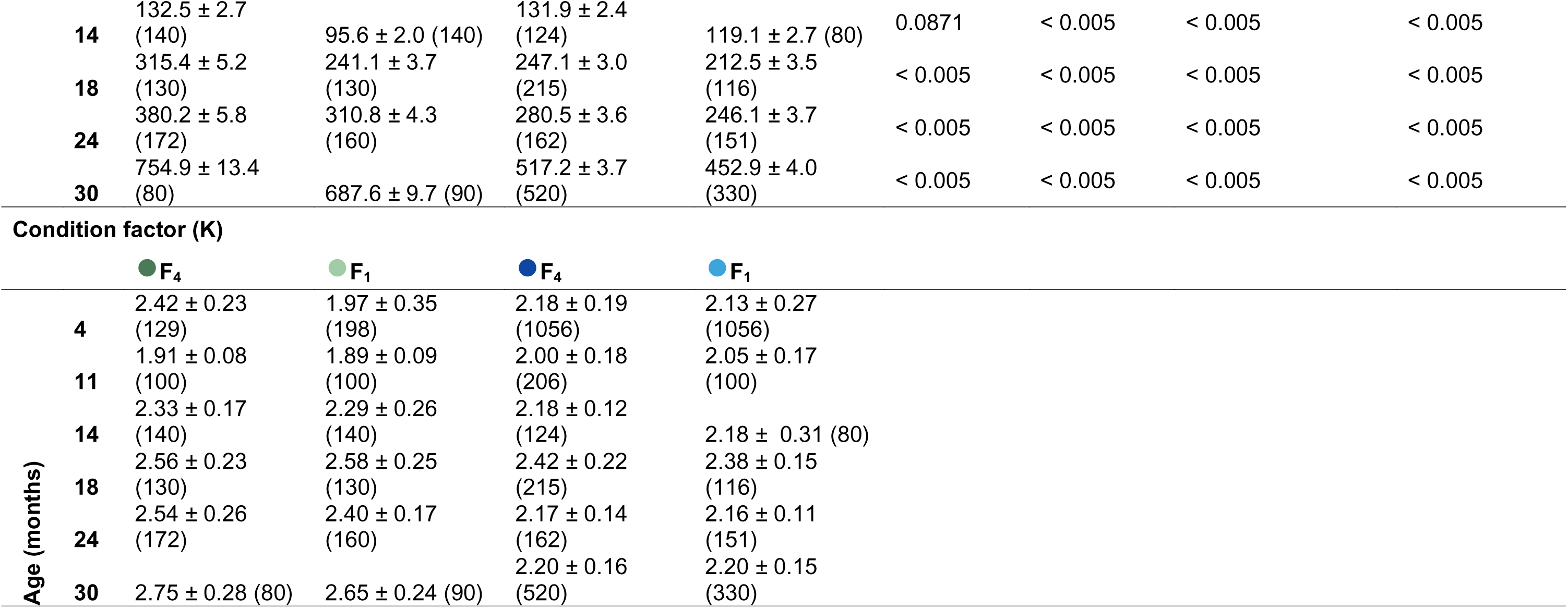
Growth of Australasian snapper (*Chrysophrys auratus*), from 4 to 30 months of age, for land-based and ocean-based F_4_ and F_1_ cohorts. The table shows the age in months down the side, then growth data for fork length (*L_F_*, mm, mean ± SE (n)). Additionally, this table shows the statistical results of a Welch’s t-test with *p*-values for within each generation across the production systems, and between generations within the production systems. This is repeated below for body weight (BW, g). Condition factor (K) is displayed at the bottom of the table, for statistics refer to the initial traits (*L_F_* and BW) prior to calculation of K.

| | | Land-based | | Ocean-based | | $p$-values (Welch t-test) | | System | |
| --- | --- | --- | --- | --- | --- | --- | --- | --- | --- |
| | | Fork length ($L_F$, mm) | | | | Generation | | | |
|  |  | ● F<sub>4</sub> | ● F<sub>1</sub> | ● F<sub>4</sub> | ● F<sub>1</sub> | F<sub>4</sub> | F<sub>1</sub> | Land-based | Ocean-based |
| Age (months) | 4 | 99 ± 1 (129) | 74 ± 1 (198) | 105 ± 0 (1056) | 93 ± 0 (1056) | < 0.005 | < 0.005 | < 0.005 | < 0.005 |
| | 11 | 144 ± 1 (100) | 131 ± 1 (100) | 152 ± 1 (206) | 143 ± 1 (100) | < 0.005 | < 0.005 | < 0.005 | < 0.005 |
| | 14 | 177 ± 1 (140) | 161 ± 1 (140) | 181 ± 1 (124) | 176 ± 1 (80) | < 0.005 | < 0.005 | < 0.005 | < 0.005 |
| | 18 | 230 ± 1 (130) | 210 ± 1 (130) | 216 ± 1 (215) | 207 ± 1 (116) | < 0.005 | 0.04248 | < 0.005 | < 0.005 |
| | 24 | 246 ± 1 (172) | 234 ± 1 (160) | 234 ± 1 (162) | 224 ± 1 (151) | < 0.005 | < 0.005 | < 0.005 | < 0.005 |
| | 30 | 301 ± 2 (80) | 296 ± 1 (90) | 286 ± 1 (520) | 273 ± 1 (330) | < 0.005 | < 0.005 | 0.01469 | < 0.005 |
|  |  | Body weight (BW, g) |  |  |  | Generation |  | System |  |
|  |  | ● F<sub>4</sub> | ● F<sub>1</sub> | ● F<sub>4</sub> | ● F<sub>1</sub> | F<sub>4</sub> | F<sub>1</sub> | Land-based | Ocean-based |
| Age (month) | 4 | 24.2 ± 0.5 (129) | 9.7 ± 0.5 (198) | 25.4 ± 0.2 (1056) | 17.9 ± 0.2 (1056) | 0.02425 | < 0.005 | < 0.005 | < 0.005 |
| | 11 | 61.8 ± 1.5 (100) | 46.8 ± 1.4 (100) | 63.1 ± 2.5 (21) | 52.7 ± 1.9 (22) | 0.6731 | 0.01274 | < 0.005 | < 0.005 |

|  |  |  |  |  |  |  |  |  |
| --- | --- | --- | --- | --- | --- | --- | --- | --- |
| 14 | 132.5 ± 2.7<br>(140) | 95.6 ± 2.0 (140) | 131.9 ± 2.4<br>(124) | 119.1 ± 2.7 (80) | 0.0871 | < 0.005 | < 0.005 | < 0.005 |
| 18 | 315.4 ± 5.2<br>(130) | 241.1 ± 3.7<br>(130) | 247.1 ± 3.0<br>(215) | 212.5 ± 3.5<br>(116) | < 0.005 | < 0.005 | < 0.005 | < 0.005 |
| 24 | 380.2 ± 5.8<br>(172) | 310.8 ± 4.3<br>(160) | 280.5 ± 3.6<br>(162) | 246.1 ± 3.7<br>(151) | < 0.005 | < 0.005 | < 0.005 | < 0.005 |
| 30 | 754.9 ± 13.4<br>(80) | 687.6 ± 9.7 (90) | 517.2 ± 3.7<br>(520) | 452.9 ± 4.0<br>(330) | < 0.005 | < 0.005 | < 0.005 | < 0.005 |

Condition factor (K)
|  | ● F<sub>4</sub> | ● F<sub>1</sub> | ● F<sub>4</sub> | ● F<sub>1</sub> |
| --- | --- | --- | --- | --- |
| Age (months) |  |  |  |  |
| 4 | 2.42 ± 0.23<br>(129) | 1.97 ± 0.35<br>(198) | 2.18 ± 0.19<br>(1056) | 2.13 ± 0.27<br>(1056) |
| 11 | 1.91 ± 0.08<br>(100) | 1.89 ± 0.09<br>(100) | 2.00 ± 0.18<br>(206) | 2.05 ± 0.17<br>(100) |
| 14 | 2.33 ± 0.17<br>(140) | 2.29 ± 0.26<br>(140) | 2.18 ± 0.12<br>(124) | 2.18 ± 0.31 (80) |
| 18 | 2.56 ± 0.23<br>(130) | 2.58 ± 0.25<br>(130) | 2.42 ± 0.22<br>(215) | 2.38 ± 0.15<br>(116) |
| 24 | 2.54 ± 0.26<br>(172) | 2.40 ± 0.17<br>(160) | 2.17 ± 0.14<br>(162) | 2.16 ± 0.11<br>(151) |
| 30 | 2.75 ± 0.28 (80) | 2.65 ± 0.24 (90) | 2.20 ± 0.16<br>(520) | 2.20 ± 0.15<br>(330) |

Individuals from the ocean-based cohorts and their associated *L_F_* and BW were able to be linked through time (start versus end time) using a biometric identification (bio-ID) derived from the captured images (Figure 3). The bio-ID uses a tri-hash method described by Arzoumanian et al. (2005) and applied previously to snapper by Samuels et al. (2024).

**Figure 3.**
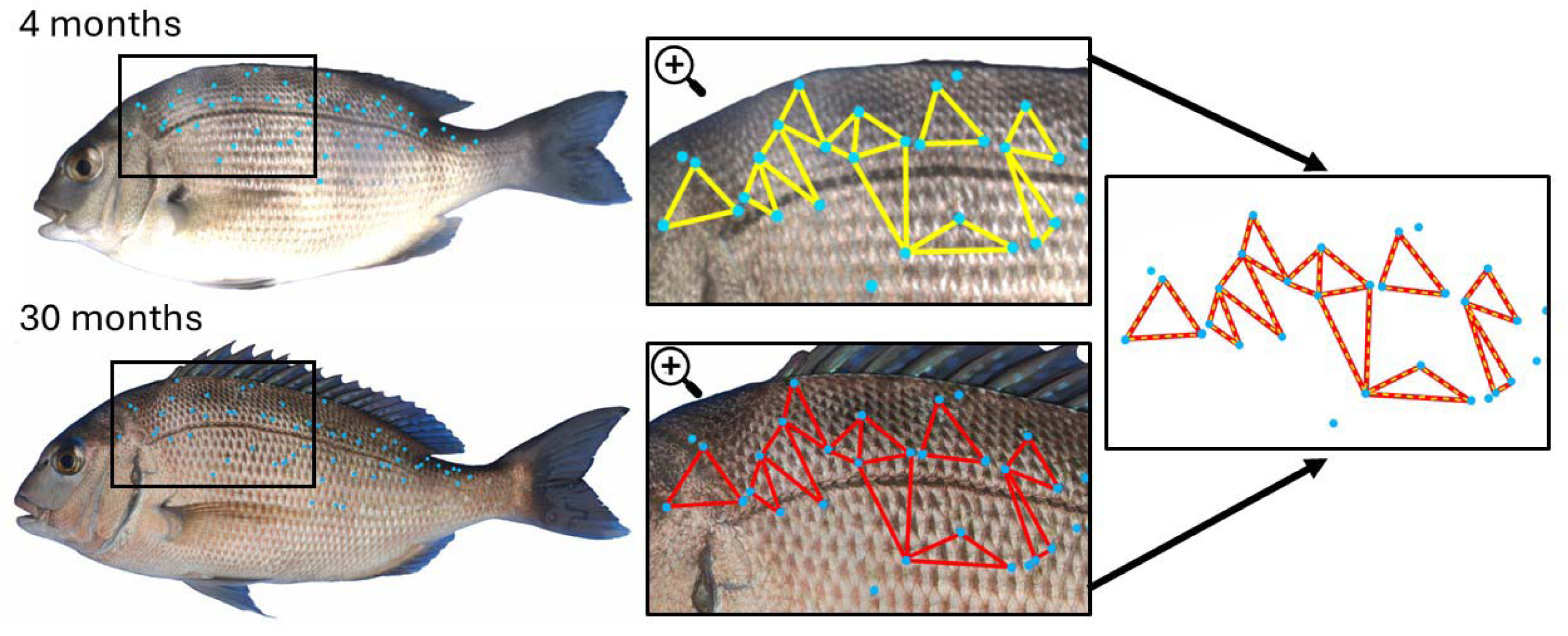
Illustration of biometric identification (bio-ID) of Australasian snapper (*Chrysophrys auratus*). The figure shows a benchtop image of the left side of the same individual at the start (4 months old) and end (30 months old) of the study. The similarities in the triangle patterns between the two time points are compared based on the methods described by Arzoumanian et al. (2005).

At 18 months of age, halfway to maturation (36 months), a sub-sample (*n = 30*) from each of the land-based F_4_ and F_1_ cohorts were euthanized for internal organ sampling to assess their condition. This sampling was restricted to the land-based cohorts because of the accessibility to freshly euthanized snapper. Data collection focused on BW (g) as well as the weights of internal organs (heart, liver, visceral fat, gastrointestinal (GI) tract, and gonads) (Figure 4).

**Figure 4.**
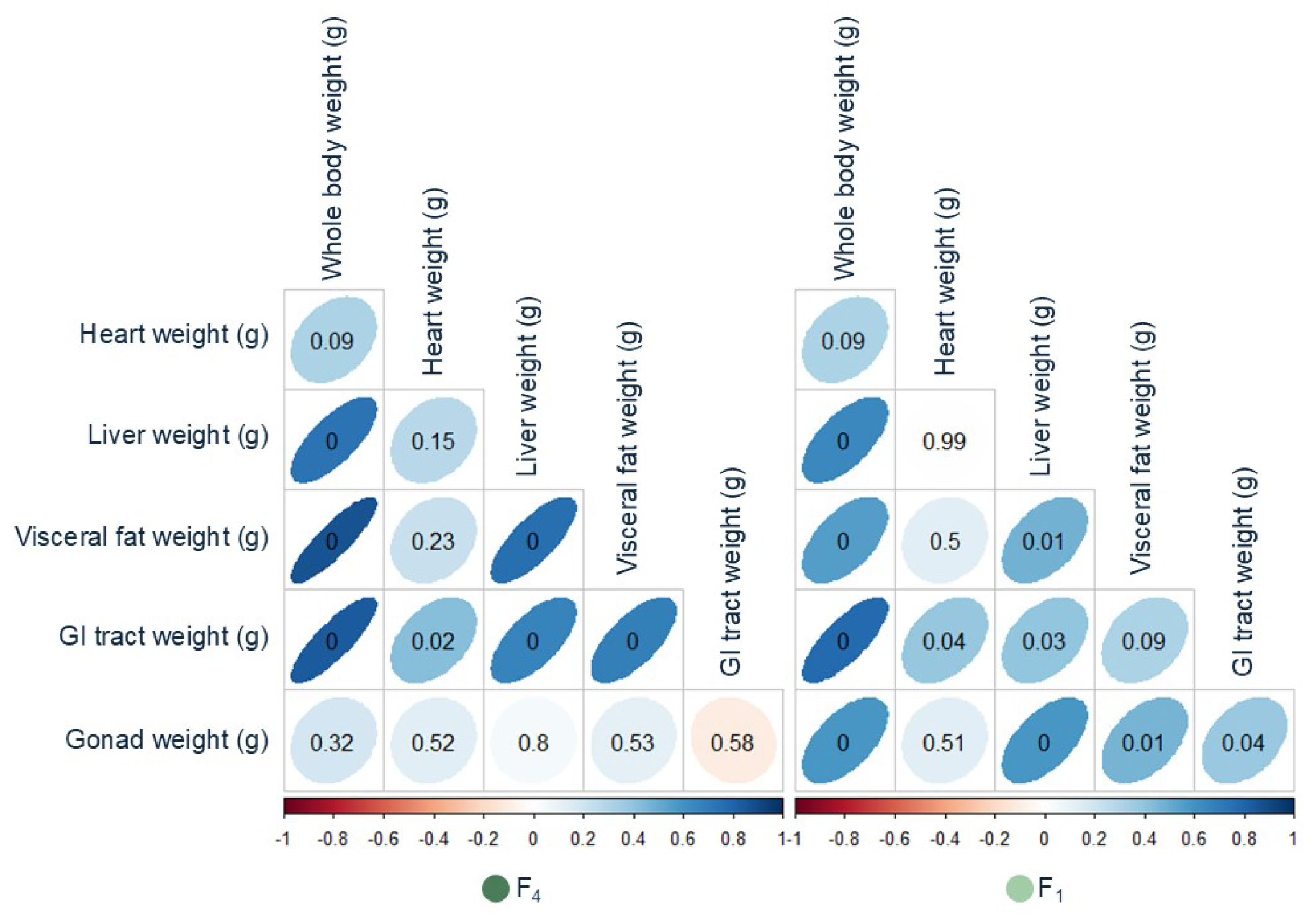
A correlation matrix between body traits for individual Australasian snapper (*Chrysophrys auratus*), including whole weight, visceral fat weight, and organ weights. Ellipse size, direction, and shade reflects R^2^ value, and the numbers in each ellipse represent the *p*-value of each correlation.

### Data analyses

Differences in the temperature profile between production sites were investigated using time-resolved plots, as well as plots of cumulative degree-days (DD, calculated by summing daily temperature), and comparison of growing degree-days (GDD, Uphoff et al. 2013) calculated using 14°C as a base temperature below which feed intake and growth effectively ceases. Where temperature data were missing at the sea pen site because of a faulty sensor and data collection device, the temperatures were linearly interpolated between the adjacent direct measurements to allow for calculation of DD and GDD.

To determine whether *L_F_* and BW improvements were statistically significant, population means were compared at each sampling event using a Welch’s t-test (Table 1), with generation and production systems as independent variables. The data were then used to calculate (a) Condition Factor (K), using the Fulton formula (K=100*BW/FL^3), where BW is body weight in grams and *L_F_* is fork length in centimetres, and (b) Coefficient of Variation (CV; CV=(mean/stdev) × 100) for the *L_F_*(mm) and body weight (g) of each cohort across all sampling events. Additionally, growth data were also used to calculate specific growth rate (SGR) (Crane et al. 2020). Temperature records for both production sites were used to calculate the thermal growth coefficient (TGC, %) (Jobling 2003).

The relationships between the lethally sampled internal traits (organ weights) and commonly assessed non-lethal traits (weight) were evaluated through a covariation matrix using the ‘corrplot’ package v0.94 (Wei and Simko 2024) in R (R Core Team 2013). Survival was calculated at the end of the study. In the land-based cohorts, survival was calculated by summing monthly mortality rates, while in the ocean-based, survival was calculated at the end of the study by comparing the stocking and retrieved numbers of the F_4_ and F_1_ snapper. To represent survival at the individual level in the sea pen, snapper IDs assigned using bio-ID were imported into R and arranged in a swarm plot based on their BW at 4 months of age.

Subsequently, BW data were highlighted in blue if the individual survived, while non-surviving individuals were highlighted in grey. Survival probability, based on fish BW at 4 months of age, was modelled using a logistic regression in tidymodels (Kuhn and Wickham 2020).

To determine whether growth rates within a population could be predicted early in the life cycle, the weight at stocking (4 months of age) was plotted against the weight at the conclusion of the study (30 months of age) and fitted with a linear regression. Additionally, all remaining individuals were ranked based on their weight at stocking and conclusion and connected between the two time points.

## Results

### Temperature comparison of on-growing sites

The temperature profile of each on-growing site differed in terms of the mean daily temperature (16.6°C for the land-based system versus 15.8°C for the ocean-based system) and the summer maximum and winter minimum temperatures, with the ocean-based system generally exhibiting a lower amplitude seasonal temperature variation (Figure 1B) than the land-based system. Using 14°C as a lower threshold for growth, the GDD metric for the land-based system was 2433 compared with 1792 for the ocean-based system (a 26% difference) over the course of the trial.

### Growth traits

Ocean-based F_4_ and F_1_ cohorts were both longer (*L_F_*) and heavier (BW) on average for the first 10 months (4 to 14 months old) of the comparison (*p<0.05*, Table 1). From this moment until the end of the study, the inverse was true, with both land-based generations growing faster and presenting significantly higher *L_F_* and BW at the end of the comparison (30 months old) (*p<0.05*, Table 1). In terms of BW at 30 months the ocean-based F_4_ and F_1_ populations were 68% and 66% of the BW recorded in the land-based cohorts, respectively.

Comparisons of F_4_ and F_1_ cohorts showed clear trends of improved growth performance in the elite strain, irrespectively of the production system. In terms of *L_F_*, F_4_ snapper already showed significant growth improvement at 4 months of age when the populations were divided into their on-growing environments, with 33.8% and 12.9% *L_F_* improvement over the F_1_ in the land-based and ocean-based cohorts, respectively (*p<0.05*, Figure 2A and Table 1). The F_4_ cohort continued to be larger than the F_1_ cohort in the land-based system, with 9.9%, 9.9%, 9.5%, 5.1% and 1.7% improvements at 11, 14, 18, 24 and 30 months respectively. Fork length improvements were also recorded in the ocean-based F_4_ cohort, with 6.3%, 2.8%, 4.3%, 4.5% and 4.8% trait improvement at 11, 14, 18, 24 and 30 months respectively (*p<0.05*, Figure 2A and Table 1). The amplitude of *L_F_* improvement in the ocean-based cohorts, however, were not as high as those observed in the land-based cohorts. In terms of BW, improvements between generations were consistently greater than those recorded for *L_F_* (Figure 2B and Table 1), with F_4_ cohorts showing significantly higher BW than F_1_ cohorts across all sampling points (*p<0.05,* Supplementary Table 2), irrespective of production system. In the land-based system, the BW of the F_4_ cohort was 149.5%, 32.1%, 38.6%, 30.8%, 22.3%, and 9.8% higher at 11, 14, 18, 24, and 30 months, respectively, than the BW of the F_1_ cohort (*p<0.05*, Table 1). The same trend was seen in the ocean-based cohorts, where the F_4_ cohort showed 41.9%, 19.7%, 10.7% 16.3%, 14.0%, and 14.2% greater BW than the F_1_ cohort at 11, 14, 18, 24, and 30 months, respectively (*p<0.05*, Figure 2B and Table 1). As seen in *L_F_*, these BW gains between cohorts were slightly smaller in the ocean-based system than those documented for the land-based system. The enhancement both *L_F_* and BW between generations was also reflected in the comparison of condition factor (K, Table 1). The K of the F_4_ cohort was higher than that of the F_1_ cohort at most time points in each production system, with the exception of 18 months in the land-based system and 11 months in the sea pen, where the F_4_ K was 0.02 and 0.05 lower than that of the F_1_ cohort, respectively (Figure 2C and Table 1).

In the F_4_ cohort, a statistically significant but moderate correlation was observed between the initial and final BW (*p < 0.001*, correlation coefficient = 0.562), suggesting that factors beyond initial BW influence growth trajectories. In contrast, no significant correlation was found for the F_1_ cohort (*p* = *0.166*), indicating weaker predictive power of initial BW in this group.

Figure 6A illustrates the start and end BW of the surviving individuals from the ocean-based cohorts. In the F_4_ cohort, the start and end BW were significantly correlated (*p* < *0.001*), with a R^2^ value of 0.271. For the F_1_ cohort, the start and end BW were also significantly correlated (p < *0.001*), with a R^2^ of 0.031. Figure 6B shows individual snapper rank progressions, based on BW, at 4 and 30 months for F_4_ and F_1_ cohorts at the sea pen. The lightest 10 and heaviest 10 fish in each generation cohort are highlighted in blue and red, respectively, at each time point.

The rank progressions also showed minimal correlations between the start and end of the comparison (F_4_ *p* < *0.001*, R^2^ = 0.31; F_1_ *p* < 0.05, R^2^ = 0.027). Individualised growth data over the same time span were not available for snapper at the land-based system.

The coefficients of variation (CV) for both *L_F_* and BW were broadly comparable between the F_1_ and F_4_ cohorts and between production systems (Table 2). There was a slight decrease in CV values variation over time, which was observed consistently across all sampled cohorts and production settings (Table 2). The specific growth rate (SGR) and temperature growth coefficient (TGC) showed that the F_4_ and F_1_ cohorts were generally comparable within production system.

**Table 2.**
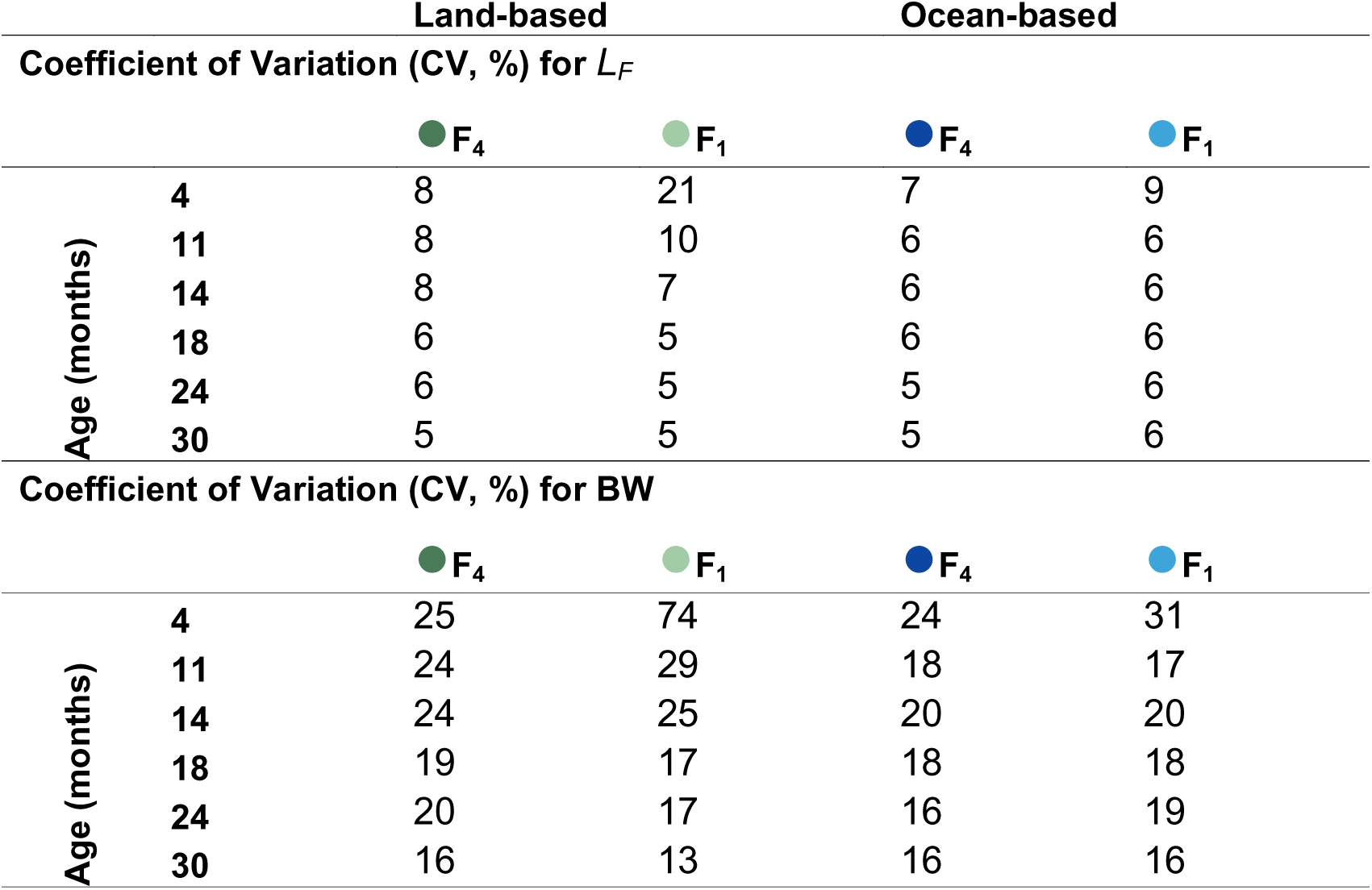
Coefficient of Variation (CV, %) for fork length (*L_F_*) and body weight (BW) of Australasian snapper (*Chrysophrys auratus*), from 4 to 30 months of age for land-based and ocean-based F_4_ and F_1_ cohorts.

|  |  | Land-based |  | Ocean-based |  |
| --- | --- | --- | --- | --- | --- |
| Coefficient of Variation (CV, %) for $L_F$ | | | | | |
| | | ● $F_4$ | ● $F_1$ | ● $F_4$ | ● $F_1$ |
| Age (months) | 4 | 8 | 21 | 7 | 9 |
|  | 11 | 8 | 10 | 6 | 6 |
|  | 14 | 8 | 7 | 6 | 6 |
|  | 18 | 6 | 5 | 6 | 6 |
|  | 24 | 6 | 5 | 5 | 6 |
|  | 30 | 5 | 5 | 5 | 6 |
| Coefficient of Variation (CV, %) for BW |  |  |  |  |  |
| | | ● $F_4$ | ● $F_1$ | ● $F_4$ | ● $F_1$ |
| Age (months) | 4 | 25 | 74 | 24 | 31 |
|  | 11 | 24 | 29 | 18 | 17 |
|  | 14 | 24 | 25 | 20 | 20 |
|  | 18 | 19 | 17 | 18 | 18 |
|  | 24 | 20 | 17 | 16 | 19 |
|  | 30 | 16 | 13 | 16 | 16 |

These values showed a marked seasonal pattern, with fish growing more slowly over winter, while faster during summer, irrespective of production system or generation. Internal traits in the F_4_ cohorts revealed significant correlations between body weight and the weights of the liver, visceral fat, and GI tract, as well as between heart and GI tract, visceral fat and GI tract, and liver with both visceral fat and GI tract (all correlations, *p* < *0.05*; Figure 4). Similarly, in the F_1_ population, significant correlations were found between body weight and the weights of the liver, visceral fat, GI tract, gonad, and heart, as well as between liver and visceral fat, GI tract, and gonad; visceral fat and gonad; and GI tract and gonad (all correlations, *p* < *0.05*; Figure 4).

### Survival

The survival of the F_1_ cohort in the land-based system from 4–30 months was 29%, and for the F_4_ cohort survival was 53%. Water temperature had a pronounced effect on mortality for both cohorts, particularly in the first year. For the F_1_ cohort the majority of mortality (76%) occurred in the first 12 months, with a pulse of mortalities occurring during the winter and autumn when waters were below approximately 15°C (Figure 5A). After this phase mortality was low and intermittent for the remainder of the trial. The F_4_ cohort showed a similar pattern of mortality although with a much lower amplitude, with comparatively fewer fish dying during the first winter (Figure 5A). The ocean-reared cohorts had similar level of survival to the land-based cohorts, with 51% survival recorded for F_4_ and 32% for F_1_. This consistency in survival rates across production systems indicates the robustness of the breeding gains, with the F_4_ cohort showing a 61-84% survival improvement over the F_1_ cohort.

**Figure 5.**
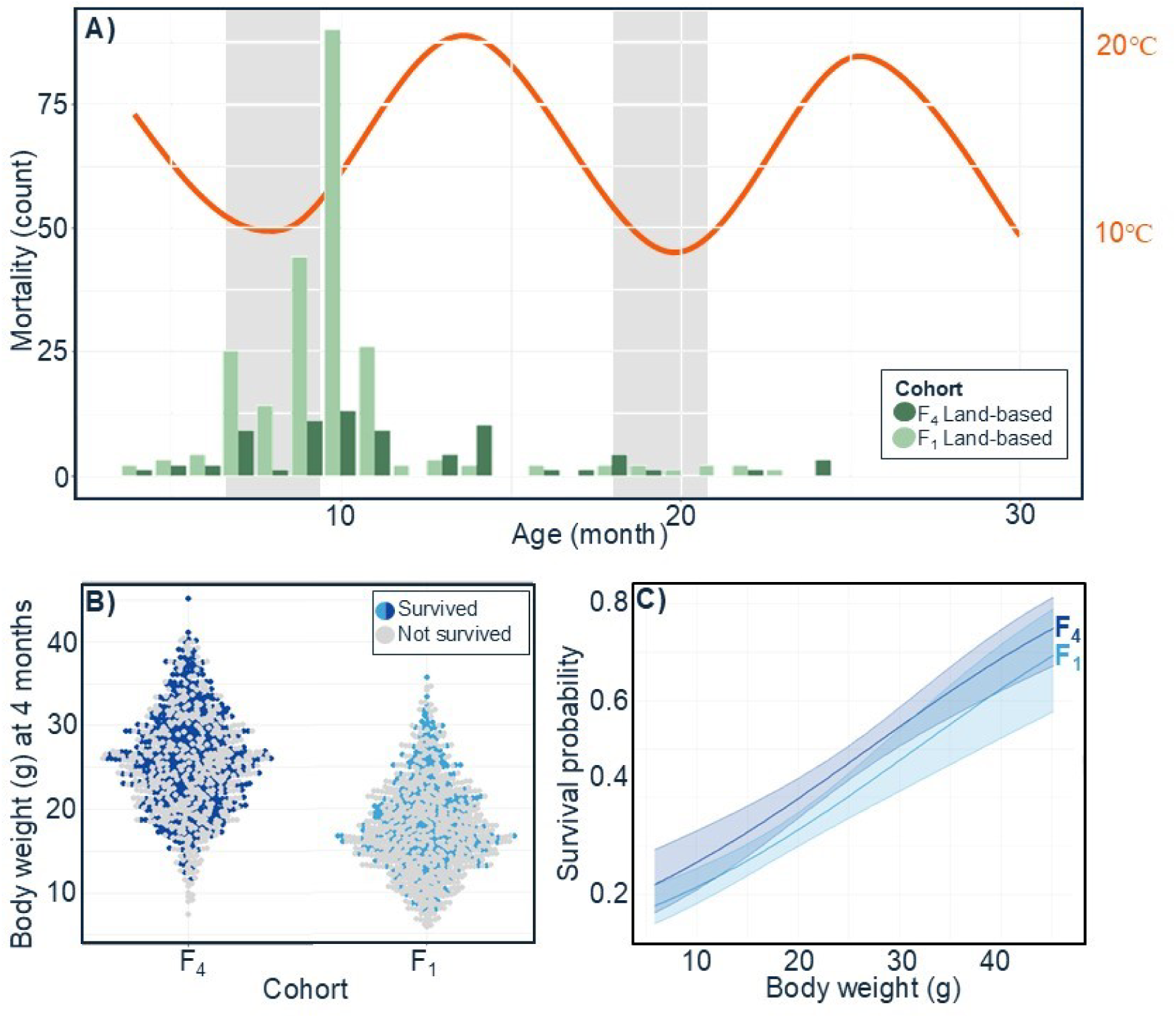
Total monthly mortality counts of land-based F_4_ and F_1_ Australasian snapper (*Chrysophrys auratus*) cohorts, from 4 to 30 months of age, coloured by cohort (F_4_ in dark green and F_1_ in light green) and the temperature profile over the same timeframe, with grey-shading areas highlighting winter months in New Zealand. Survival of ocean-based cohorts are shown in panel B, where all individual snapper are arranged by body weight at 4 months of age for both F_4_ and F_1_ cohorts. Surviving individuals at the end of the comparison (30 months) are highlighted, while those who did not survive are in grey. Panel C shows a logistic regression model of survival versus body weight, based on body weight at 4 months of age and their survival outcome at the end of the comparison. Shading indicates the 95% confidence interval for each model.

**Figure 6.**
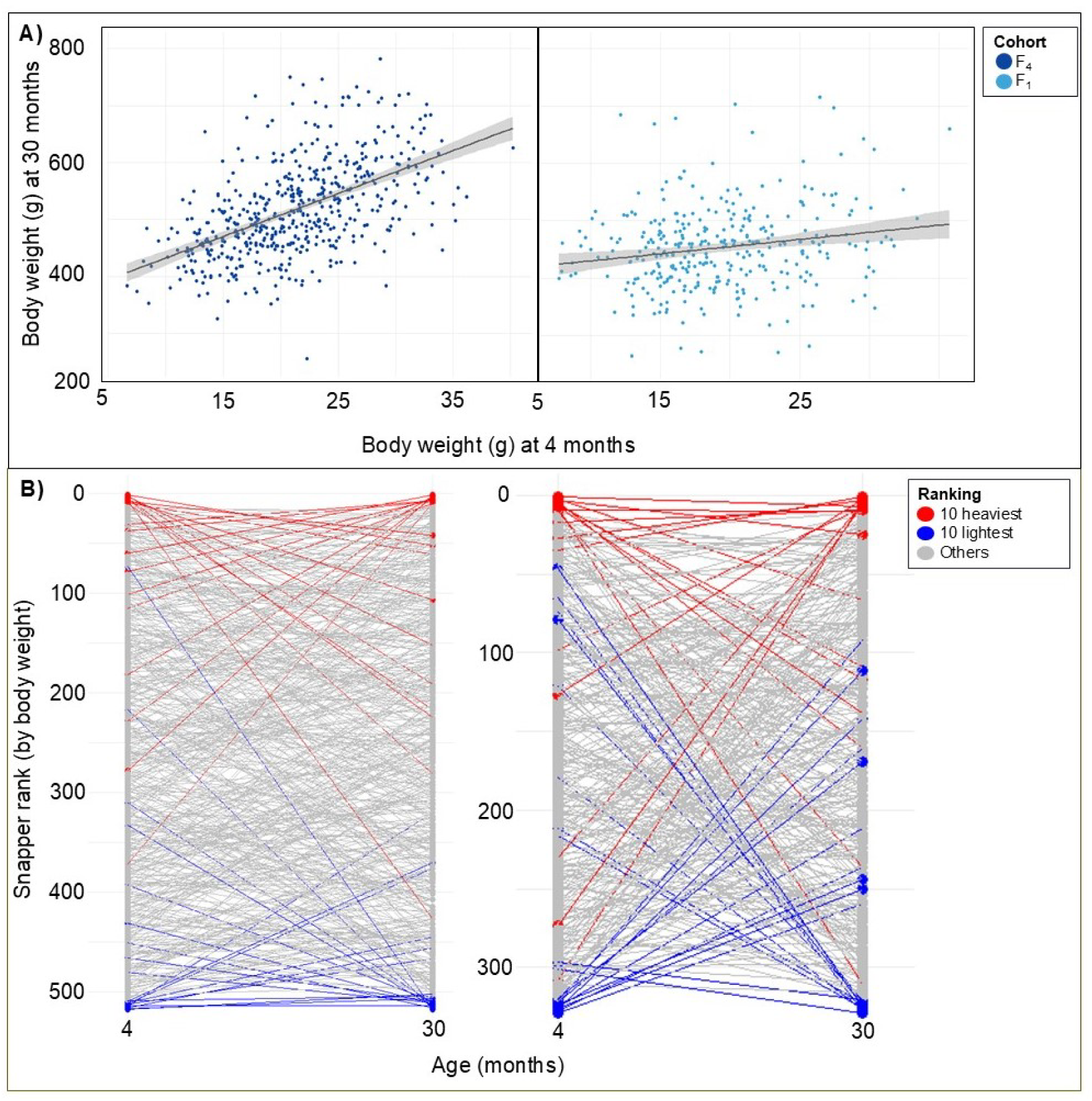
Panel A: Scatterplot showing the body weight of ocean-based grown F_4_ and F_1_ Australasian snapper (*Chrysophrys auratus*) at two time points—4 months (start) and 30 months (end) of the comparison—fitted with a linear regression line and associated 95% confidence interval to illustrate the relationship between early and later body weight. Panel B: Rank progression plot of individual snapper based on body weight at the start and end of the comparison. The lightest 10 and heaviest 10 individuals at each time point are highlighted in blue and red, respectively.

The biometric re-identification tool enabled analysis of size-specific mortality for each cohort in the ocean-based system as each surviving fish at 30 months could be linked to its corresponding weight at 4 months (Figure 5B). A logistic regression model that plotted survival probability against weight at 4 months indicated a strong size-selective survival function for both cohorts, with fish around 10 g having a 20% probability of survival to 30 months, whereas fish around 40 g had a more than 60% survival rate (Figure 5C). There was small but consistent separation in the survival function between cohorts, with the F_4_ cohort demonstrating a 5-10% elevated survival probability for any given size, however, the confidence intervals for two regressions were overlapping, meaning the difference wasn’t statistically significant at the 95% level.

## Discussion

To meet the future food needs of a growing global population and amidst warming sea temperatures, species diversification in aquaculture is essential (Lubchenco et al. 2020; Naylor et al. 2021). Indeed, the diversification of species is seen as a key adaptation strategy to enhance the resilience of the aquaculture sector against the impacts of climate change (Cai et al. 2023; Edgar et al. 2024).

However, implementing species diversification is a long-term strategy, requiring careful consideration of factors such as species selection and site suitability (Metian et al. 2020). Here, we present for the first time results of a full on-growing cycle of snapper *Chrysophrys auratus* reared in the Top of the South Island, New Zealand.

Moreover, we compare the performance of both F_4_ (bred for enhanced growth) and F_1_ (offspring of wild-caught broodstock) and generations of snapper grown in a flow-through facility (land-based) in Nelson and a sea pen in Marlborough Sounds (ocean-based).The Marlborough Sounds is a traditionally used exclusively for Chinook salmon (*Oncorhynchus tshawytscha*) farming; however, recent climate change has caused frequent and extended marine heatwaves, raising water temperatures above 18°C for extended periods (Law et al. 2018; Cook et al. 2024). This is stressful for Chinook salmon and has been linked to increased mortality rates and heighted vulnerabilities to pathogens.

Our aim in this study was to evaluate the performance of snapper in a realistic commercial aquaculture sea pen setting in the Marlborough Sounds, with the ultimate goal of determining the species potentiality as a viable alternative to salmon farming in the region. Our study revealed that the F_4_ line grew significantly faster than the unselected snapper, both in land-based and ocean-based production systems. In the land-based cohorts, the elite F_4_ cohort achieved 400 grams in approximately 740 days, and 600 grams in about 810 days. In contrast, the F_1_ cohort reached 400 grams in around 790 days and 600 grams in approximately 840 days.

In the ocean-based cohorts, the F_4_ cohort reached 400 grams in roughly 800 days, while the F_1_ cohort reached 400 grams in about 850 days. Differences in water temperature was likely the most important explanatory factor, with a substantially lower sum of growing degree days in the ocean-based site (1972 versus 2433, a 26% difference) correlating with a 32-34% lower mean body weight compared to fish from the land-based site. The difference in temperature regimes between sites despite a relatively short distance (65 km) reflects the respective coastal typographies, with the land-based Nelson site drawing water from a shallow estuarine harbour that was more influenced by air temperature than the sea pen located in the seaward reaches of the Marlborough Sounds. It is important to note that for our experiments we deliberately used a low level of size grading as we wanted to preserve phenotypic variability to understand how a wide selection of the populations performed. This contrasts with commercial settings, where higher grading would typically be used early during the nursery stage to remove underperforming and small fish, meaning the average growth rates reported in this study are a low-end estimate if one wishes to extrapolate the data to a commercial production model.

The fast growth observed in the selectively bred snapper aligns with results from selective breeding programmes in other species within the sparid family, particularly the red sea bream (*Pagrus major*) and gilthead sea bream (*Sparus aurata*). The red sea bream is primarily cultured in Japan, while the gilthead sea bream is farmed around the Mediterranean. Typically, both species are harvested between 18 and 24 months, depending on farming practices and market size (Mhalhel et al. 2023). The Aquaculture Research Institute at Kindai University (ARIKU) in Japan has been selectively breeding red sea bream since the early 1960s, achieving a harvest weight of approximately 1000 grams after around 738 days, plus an additional 48 days for further growth (Murata et al. 1996; Kato 2023). In contrast, gilthead sea bream, which has been selectively bred by various companies since the late 1980s, is harvested at smaller sizes, starting from 400 grams (Gulzari et al. 2022). Selective breeding has significantly enhanced growth rates and feed efficiency for both species, with red sea bream exhibiting a 90% weight improvement and gilthead sea bream showing growth rate improvements of 15–30% over non-selected lines (Boudry et al. 2021). Phenotypic trait correlations showed strong positive allometry between weight and length for both cohorts, indicating integrated growth and development. This scaling suggests that as snapper grow, traits like internal organ mass adapt proportionally to support physiological demands. These findings are valuable for selective breeding, as selecting for weight and length may indirectly enhance traits critical for growth and health, improving overall aquaculture performance (Klingenberg 1996; Karachle and Stergiou 2012).

Survival was also generally higher in the F_4_ line than in the unselected F_1_ cohort, with more than half the F_4_ fish surviving in both land-based and ocean-based systems, while only around 30% of the F_1_ cohort survived in these environments.

Survival of the F_1_ cohort was slightly better in the ocean-based environment, suggesting that the sea pen may more closely resemble the natural conditions to which the F_1_ snapper, produced from wild-spawned parents, are adapted. These wild snapper were not selected for optimal performance in captive settings, unlike the breeding line, which has been exposed for 20 years to these conditions (Baesjou and Wellenreuther 2021; Moran et al. 2023; Samuels et al. 2024). The consistency in survival rates for each generation, regardless of the production system, strengthens the reliability of these survival results.

The use of the two production systems to study distinct aspects of survival helped elucidate important biological traits of this species that impact aquaculture production strategies for the South Island. The land-based system enabled easy monitoring of fish populations and characterisation of survival patterns throughout seasons, while the passive biometric tagging of fish in the ocean-based system enabled a longitudinal analysis of juvenile stocking size and survival in a commercial sea pen setting. The land-based observations indicated that survival to harvest is highly impacted by the first winter a juvenile experiences, and the ocean-based data revealed that there is a strong size-selective survival effect. Putting these two pieces of information together, we hypothesize that for snapper to have a high survival rate (>90%) for aquaculture production it is critical that juveniles are at least 60 g before they encounter the cooler winter temperatures that cause feeding to cease at around 14°C. We assume that this minimum weight is required to support physiological processes that continue for the 2-3 months of winter when feed intake is low or zero. Tools aquaculturists have to achieve this size and timing specification includes inducing early spawning to ensure snapper have time to reach the target weight before winter and size grading prior to stocking in net pens. These findings for snapper mirror those observed for yellowtail kingfish, where Australian aquaculture researchers found juveniles of this species highly susceptible to low winter temperatures, and larger individuals over 44 g were more robust to the cool winter waters of the Spencer Gulf when stocked in net pens (Booth M, Allan, et al. 2008).

Our study design and findings on growth and survival are significant, as they address the challenges of evaluating production-related breeding gains for a new candidate species for aquaculture, particularly in terms of cost-effectiveness and real-world applicability. A key aspect of our approach was the ability to mix cohorts within the same pen and later separate them using phenotypic metrics. This cost-efficient strategy minimizes variability in growing environments, enabling more accurate cohort comparisons and providing realistic insights into comparative gains and their relationship with environmental factors. Our findings also highlight several important challenges. Performance was shown to be cohort-dependent, with the F_4_ outperforming the F_1_ cohort, as well as several indicators showing that performance is both environment- and age-dependent. Notably, we found that optimal performance varied over the course of the 2-year experiment, suggesting that grow-out strategies may need to be tailored to the life stages of fish selected for specific production systems. This has critical implications for maximizing economic gains and production efficiency. Specifically, both F_4_ and F_1_ snapper cohorts exhibited greater growth gains in the sea pen during the initial 10 months of the comparison. However, over the subsequent 16 months both cohorts grew faster in the land-based system.

Future research should prioritize the continued use of selective breeding as a powerful tool to enhance the commercial viability of snapper aquaculture, with a view to enhance not only growth rates but also resilience to cold water temperatures.

While significant progress has been made, particularly in earlier generations, such as the feed efficiency improvements observed in the F_3_ cohort compared with the F_1_ cohort (Moran et al. 2023), there remains a critical gap in understanding growth and survival dynamics during the grow-out phase, particularly in coastal and offshore net pen systems. Most studies in New Zealand and Australia to date have been conducted in land-based systems, leaving a lack of data on performance in sea pen environments. A key consideration for any successful commercial breeding programme is the need to replicate real-world farming conditions during the selection process. By doing so, animals evaluated and selected are more likely to express desired traits consistently under commercial operations, such as improved growth and survival. This alignment is critical not only for ensuring that breeding gains translate into tangible benefits for producers but also for supporting the long-term resilience and sustainability of the breeding line.

## Conclusions

Aquaculture species diversification is becoming increasingly urgent worldwide, particularly in New Zealand, as climate change-induced marine heatwaves intensify and lengthen (Law et al. 2018; Laufkötter et al. 2020). The Top of the South Island is especially significant in this context, as it already hosts established coastal aquaculture infrastructure for Chinook salmon (*Oncorhynchus tshawytscha*). However, rising water temperatures—consistently exceeding 18°C in recent years—have led to substantial losses of Chinook salmon in the top of the south of New Zealand, prompting interest in developing more climate-resilient aquaculture operations. Snapper emerges as a promising candidate for such diversification, thriving in water temperatures between 18 and 24°C (Bowering et al. 2023; Collins et al. 2024), a range that fosters improved growth rates. Our results demonstrated that the selectively bred aquaculture line of snapper reached harvestable sizes within two years of being placed in the sea pen, showing better survival rates than unselected snapper. These growth and survival performance metrics are expected to improve significantly if a graded cohort approach is applied, selecting only the top-performing snapper based on body weight, which was not implemented in this study. With such selective grading, coupled with twice-daily optimal feeding, production gains would probably surpass those reported here.

Moreover, strategies such as advance broodstock spawning through photothermal manipulation have the potential to anticipate the transference of fingerlings into the sea pen to spring or early summer. This strategy can significantly boost early cohort growth, leading to robust and bigger fish to undergo first winter, probably boosting survival. Furthermore, nutrition strategies as those suggested for Nile tilapia (Nobrega et al. 2020), can boost health, survival and performance of warm-water species during the colder months. Taken together, these initial findings provide valuable insights into the potential of snapper as a climate-resilient species for aquaculture in this region, offering an alternative to traditional salmon farming. Further research is required to optimize grading protocols, production, feeding and nutrition strategies to maximize growth rates, survival and improve economic returns.

## Acknowledgements and Funding details

We thank all the staff of The New Zealand Institute for Plant and Food Research Limited who were involved in breeding and rearing the snapper populations that formed part of this breeding programme. This project was supported by a New Zealand Ministry of Business, Innovation & Employment Endeavour Fund (C11X1603) ‘Accelerated breeding for enhanced seafood production’ to MW and a Technology Development Fund from The New Zealand Institute for Plant and Food Research Limited to MW.

## Data availability

Raw data sets are available on request from the corresponding author.

## Supplementary Materials

### Tables

**S. Table 1.**
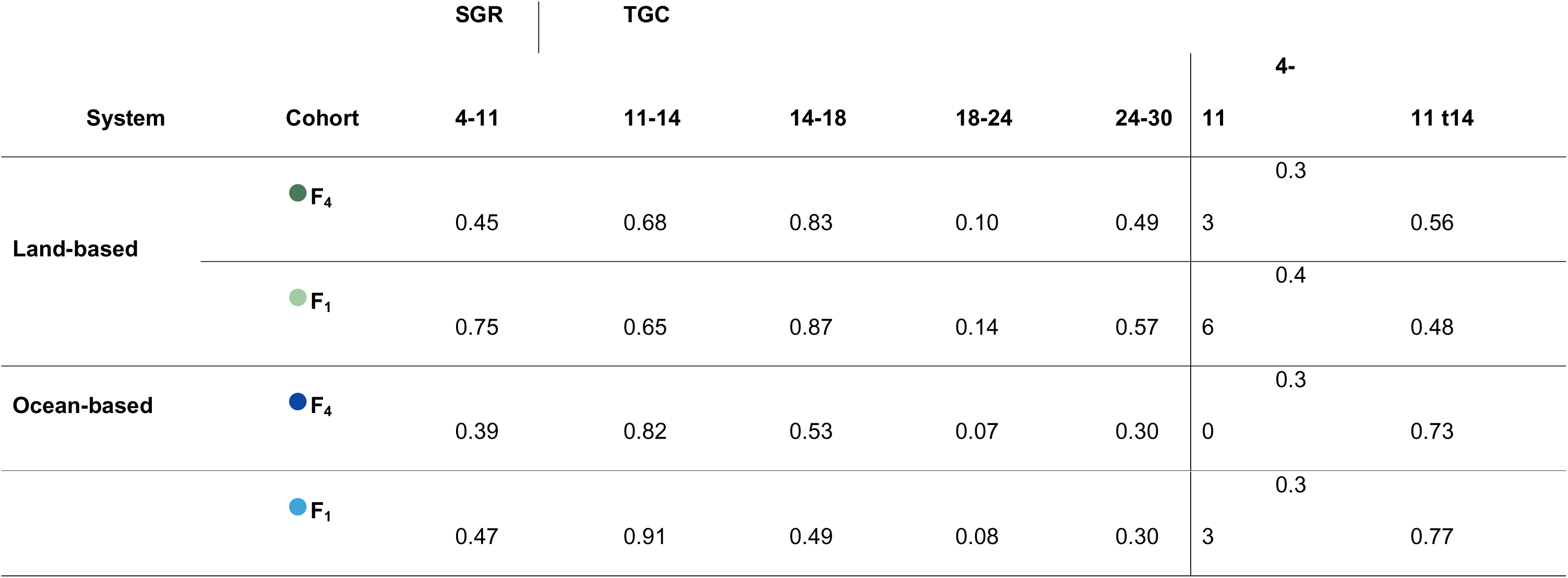
Specific Growth Rate (SGR, % day^-1^) and Thermal Growth Coefficient (TGC% day^-1^), based on body weight of Australasian snapper (*Chrysophrys auratus*) between sampling events from 4 to 30 months of age, for land-based and ocean-based F_4_ and F_1_ cohorts.

### Figures

**S Figure 1.**
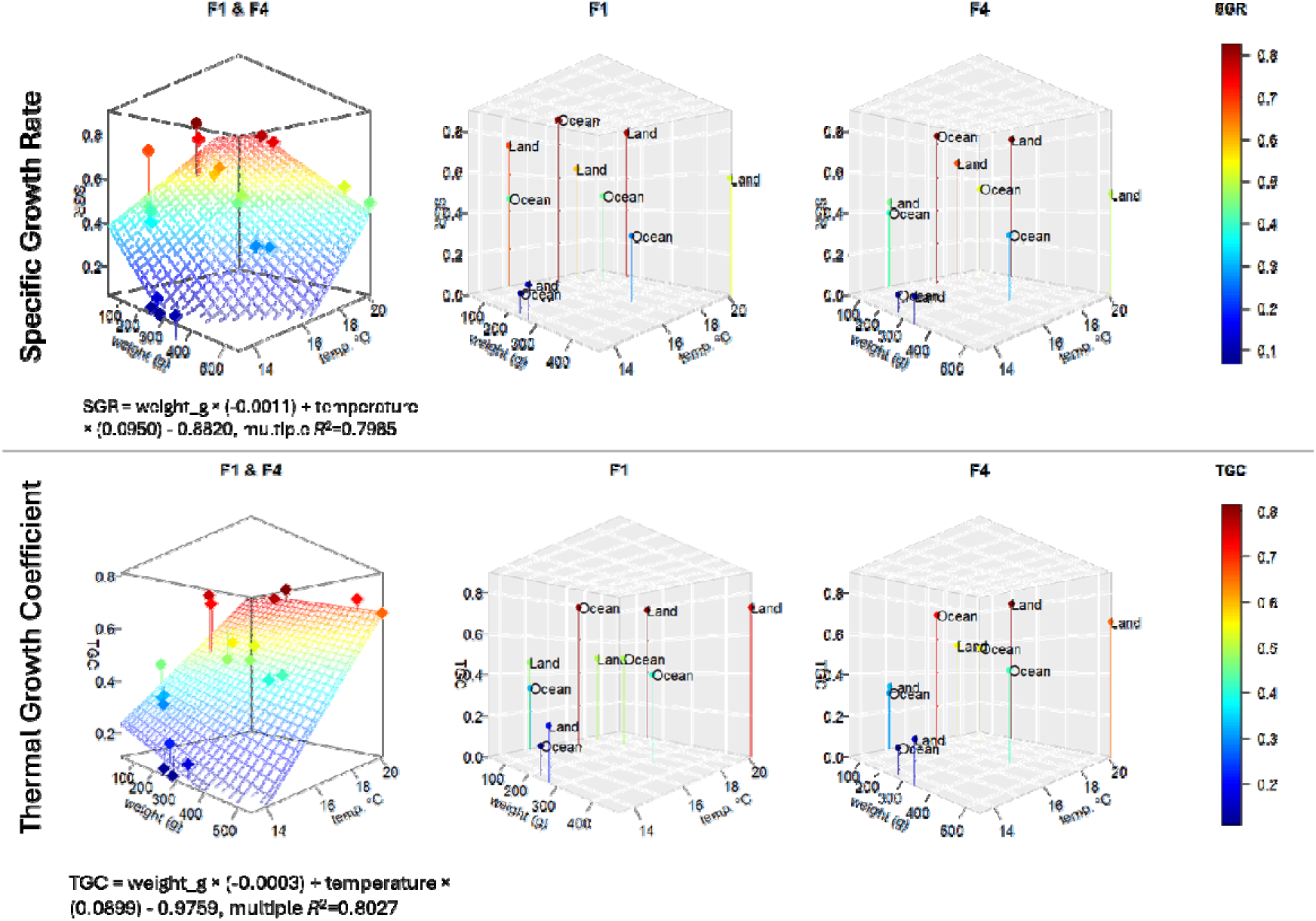
Plots of Specific Growth Rate (% body weight day^-1^) and Thermal Growth Coefficient versus body weight (geometric mean for measurement interval) of Australasian snapper (*Chrysophrys auratus*) and water temperature (mean temperature for measurement interval).

## References

Arzoumanian Z, Holmberg J, Norman B. 2005. An astronomical pattern-matching algorithm for computer-aided identification of whale sharks Rhincodon typus. J APPL ECOL. 42(6):999–1011.

Baesjou JP, Wellenreuther M. 2021. Genetic signatures of domestication selection in the Australasian snapper (*Chrysophrys auratus*). Genes. 12(11):1737.

Booth M, Allan G, Russell I, Elkins M, Bowyer J. 2008. Understanding yellowtail kingfish: sub-project 3. Nova. 100(110):120.

Booth M, Tucker B, Allan G, Fielder DS. 2008. Effect of feeding regime and fish size on weight gain, feed intake and gastric evacuation in juvenile Australian snapper Pagrus auratus. Aquaculture. 282(1-4):104–110.

Booth MA, Allan GL, Anderson AJ. 2007. Investigation of the nutritional requirements of Australian snapper Pagrus auratus (Bloch & Schneider, 1801): effects of digestible energy content on utilization of digestible protein. Aquaculture Research. 38(4):429–440.

Booth MA, Warner-Smith RJ, Allan GL, Glencross BD. 2004. Effects of dietary astaxanthin source and light manipulation on the skin colour of Australian snapper Pagrus auratus (Bloch & Schneider, 1801). Aquaculture Research. 35(5):458–464.

Boudry P, Allal F, Aslam ML, Bargelloni L, Bean TP, Brard-Fudulea S, Brieuc MSO, Calboli FCF, Gilbey J, Haffray P et al. 2021. Current status and potential of genomic selection to improve selective breeding in the main aquaculture species of International Council for the Exploration of the Sea (ICES) member countries. Aquaculture Reports. 20:100700.

Bowering LR, McArley TJ, Devaux JB, Hickey AJ, Herbert NA. 2023. Metabolic resilience of the Australasian snapper (Chrysophrys auratus) to marine heatwaves and hypoxia. Frontiers in Physiology. 14:1215442.

Boyd CE, McNevin AA, Davis RP. 2022. The contribution of fisheries and aquaculture to the global protein supply. Food Sec. 14(3):805–827.

Cai J, Chan HL, Yan X, Leung P. 2023. A global assessment of species diversification in aquaculture. Aquaculture. 576:739837.

Collins SP, Magnoni LJ, Cook DG, Black SE, Herbert NA. 2024. What temperature is best for the offshore farming of the Australasian snapper, Chrysophrys auratus? A collective examination of growth, FCR, O2 consumption and welfare. New Zeal J Mar Fresh.1–24.

Cook KM, Dunn MR, Behrens E, Pinkerton MH, Law CS, Cummings VJ. 2024. The impacts of marine heatwaves on ecosystems and fisheries in Aotearoa New Zealand. New Zeal J Mar Fresh.1–31.

Crane DP, Ogle DH, Shoup DE. 2020. Use and misuse of a common growth metric: guidance for appropriately calculating and reporting specific growth rate. Reviews in Aquaculture. 12(3):1542–1547.

Crossland J. 1981a. biology of the New Zealand snapper. Occassional publication-Fisheries Research Division.

Crossland J. 1981b. Fish eggs and larvae of the Hauraki Gulf, New Zealand. Fisheries Research Division, New Zealand Ministry of Agriculture and Fisheries.

De Boni D. 2025 Warm seas boost fish mortality - and take a bite out of next year’s profit at NZ King Salmon. The Post.

De Meester L, Vázquez-Domínguez E, Kassen R, Forest F, Bellon MR, Koskella B, Scherson RA, Colli L, Hendry AP, Crandall KA. 2024. A link between evolution and society fostering the UN sustainable development goals. Evol Appl. 17(6):e13728.

Doolan BJ, Booth MA, Jones PL, Allan GL. 2007. Effect of cage colour and light environment on the skin colour of Australian snapper Pagrus auratus (Bloch & Schneider, 1801). Aquaculture Research. 38(13):1395–1403.

The economic contribution of commercial fishing. 2022. https://deepwatergroup.org/wp-content/uploads/2022/09/BERL-2022-Commercial-Fishing-Economic-Contribution-Final-Report.pdf.

Edgar GJ, Bates AE, Krueck NC, Baker SC, Stuart-Smith RD, Brown CJ. 2024. Stock assessment models overstate sustainability of the world’s fisheries. Science. 385(6711):860–865.

Falconer L, Sparboe LO, Dale T, Hjøllo SS, Stavrakidis-Zachou O, Bergh Ø, James P, Papandroulakis N, Puvanendran V, Siikavuopio SI. 2024. Diversification of marine aquaculture in Norway under climate change. Aquaculture. 593:741350.

mFAO R. 2022. The state of world fisheries and aquaculture 2022. Towards blue transformation. Food and Agriculture Organization of the United Nations Rome, Italy. p. 266.

Ferrell D. 1993. Assessment of the fishery for snapper (Pagrus auratus) in Queensland and New South Wales. Final Report, FRDC Project 93/074. 143.

Fielder DS, Bardsley WJ, Allan GL. 2001. Survival and growth of Australian snapper, Pagrus auratus, in saline groundwater from inland New South Wales, Australia. Aquaculture. 201(1-2):73–90.

Folkvord A, Fiksen Ø, Høie H, Johannessen A, Otterlei E, Vollset KW. 2009. What can size distributions within cohorts tell us about ecological processes in fish larvae? Scientia Marina. 73(S1):119–130.

Francis MP. 1996. Geographic distribution of marine reef fishes in the New Zealand region. New Zeal J Mar Fresh. 30:30–55.

Francis MP. 2001. Coastal fishes of New Zealand. An identification guide. 3rd edn. ed. Vol. 3rd edn. Auckland: Reed Books.

Garrido S, Ben-Hamadou R, Santos AMP, Ferreira S, Teodósio M, Cotano U, Irigoien X, Peck MA, Saiz E, Re P. 2015. Born small, die young: Intrinsic, size-selective mortality in marine larval fish. Scientific reports. 5(1):17065.

Graham D. 1953. A treasury of New Zealand fish. Wellington (New Zealand): AH & AW Reed.

Gulzari B, Komen H, Nammula VR, Bastiaansen JWM. 2022. Genetic parameters and genotype by environment interaction for production traits and organ weights of gilthead seabream (Sparus aurata) reared in sea cages. Aquaculture. 548:737555.

Jobling M. 2003. The thermal growth coefficient (TGC) model of fish growth: a cautionary note. Aquaculture Research. 34(7):581–584.

Karachle PK, Stergiou KI. 2012. Morphometrics and allometry in fishes. morphometrics.65–86.

Kato K. 2023. Breeding studies on red sea bream Pagrus major: mass selection to genome editing. Fisheries Science. 89(2):103–119.

Klingenberg CP. 1996. Multivariate allometry. Advances in morphometrics. Springer; p. 23–49.

Kuhn M, Wickham H. 2020. Tidymodels: a collection of packages for modeling and machine learning using tidyverse principles. [accessed]. https://www.tidymodels.org.

Laufkötter C, Zscheischler J, Frölicher TL. 2020. High-impact marine heatwaves attributable to human-induced global warming. Science. 369(6511):1621–1625.

Law CS, Rickard GJ, Mikaloff-Fletcher SE, Pinkerton MH, Behrens E, Chiswell SM, Currie K. 2018. Climate change projections for the surface ocean around New Zealand. New Zeal J Mar Fresh. 52(3):309–335.

Le Pape O, Bonhommeau S. 2015. The food limitation hypothesis for juvenile marine fish. Fish Fish. 16(3):373–398.

Leach F. 2007. Fishing in Pre-European New Zealand. NZ ASA. 15:469–471.

Lubchenco J, Haugan PM, Pangestu ME. 2020. Five priorities for a sustainable ocean economy. Nature. 588(7836):30–32.

Meng H, Hayashida H, Norazmi-Lokman NH, Strutton PG. 2022. Benefits and detrimental effects of ocean warming for Tasmanian salmon aquaculture. Continental Shelf Research. 246:104829.

Metian M, Troell M, Christensen V, Steenbeek J, Pouil S. 2020. Mapping diversity of species in global aquaculture. Rev Aquac. 12(2):1090–1100.

Mhalhel K, Levanti M, Abbate F, Laurà R, Guerrera MC, Aragona M, Porcino C, Briglia M, Germanà A, Montalbano G. 2023. Review on gilthead seabream (*Sparus aurata*) aquaculture: life cycle, growth, aquaculture practices and challenges. Journal of Marine Science and Engineering. 11(10):2008.

Moran D, Schleyken J, Flammensbeck C, Fantham W, Ashton D, Wellenreuther M. 2023. Enhanced survival and growth in the selectively bred *Chrysophrys auratus* (Australasian snapper, tāmure). Aquaculture. 563 (1)

Murata O, Harada T, Miyashita S, Izumi K-i, Maeda S, Kato K, Kumai H. 1996. Selective breeding for growth in red sea bream. Fisheries science. 62(6):845–849.

Naylor RL, Hardy RW, Buschmann AH, Bush SR, Cao L, Klinger DH, Little DC, Lubchenco J, Shumway SE, Troell M. 2021. A 20-year retrospective review of global aquaculture. Nature. 591(7851):551–563.

Nobrega RO, Banze JF, Batista RO, Fracalossi DM. 2020. Improving winter production of Nile tilapia: What can be done? Aquaculture Reports. 18:100453.

Papa Y, Oosting T, Valenza-Troubat N, Wellenreuther M, Ritchie PA. 2020. Genetic stock structure of New Zealand fish and the use of genomics in fisheries management: an overview and outlook. NZ J Zool. 48:1–31.

Parsons D, Sim-Smith C, Cryer M, Francis M, Hartill B, Jones E, Le Port A, Lowe M, McKenzie J, Morrison M. 2014. Snapper (*Chrysophrys auratus*): a review of life history and key vulnerabilities in New Zealand. New Zeal J Mar Fresh. 48:256–283.

Peres HA, Robert D, Mainguy J, Sirois P. 2022. Interannual variability in size-selective winter mortality of young-of-the-year striped bass. ICES J Mar Sci. 79(5):1614–1623.

R Core Team. 2013. A language and environment for statistical computing. R Foundation for Statistical Computing. http://www.R-project.org/. Vienna, Austria.

Rosenberg A, Haugen A. 1982. Individual growth and size-selective mortality of larval turbot (Scophthalmus maximus) reared in enclosures. Mar Biol. 72:73–77.

Samuels G, Hegarty L, Fantham W, Ashton D, Blommaert J, Wylie MJ, Moran D, Wellenreuther M. 2024. Generational breeding gains in a new species for aquaculture, the Australasian snapper (*Chrysophrys auratus*). Aquaculture.(586):740782.

Smith MD, Roheim CA, Crowder LB, Halpern BS, Turnipseed M, Anderson JL, Asche F, Bourillón L, Guttormsen AG, Khan A. 2010. Sustainability and global seafood. Science. 327(5967):784–786.

Sogard SM. 1997. Size-selective mortality in the juvenile stage of teleost fishes: a review. Bull Mar Sci. 60(3):1129–1157.

Teletchea F. 2021. Fish domestication in aquaculture: 10 unanswered questions. Animal Frontiers. 11(3):87–91.

United Nations. 2015. Sustainable development goals and the ocean. [accessed 2021 February]. https://en.unesco.org/sustainabledevelopmentgoalsandtheocean.

Uphoff CS, Schoenebeck CW, Hoback WW, Koupal KD, Pope KL. 2013. Degree-day accumulation influences annual variability in growth of age-0 walleye. Fish Res. 147:394–398.

Wei T, Simko V. 2024. R package ‘corrplot’: Visualization of a Correlation Matrix (Version 0.94). [accessed]. https://github.com/taiyun/corrplot

Wellenreuther M, Le Luyer J, Cook D, Ritchie PA, Bernatchez L. 2019. Domestication and temperature modulate gene expression signatures and growth in the Australasian snapper *Chrysophrys auratus*. G3: Genes, Genomes, Genetics. 9(1):105–116.

Yadav NK, Patel AB, Singh SK, Mehta NK, Anand V, Lal J, Dekari D, Devi NC. 2024. Climate change effects on aquaculture production and its sustainable management through climate-resilient adaptation strategies: a review. Environmental Science and Pollution Research. 31(22):31731–31751.

Ytteborg E, Falconer L, Krasnov A, Johansen L-H, Timmerhaus G, Johansson GS, Afanasyev S, Høst V, Hjøllo SS, Hansen ØJ. 2023. Climate change with increasing seawater temperature will challenge the health of farmed Atlantic Cod (Gadus morhua L.). Front Mar Sci. 10:1232580.

